# A critical initialization for biological neural networks

**DOI:** 10.1101/2025.01.10.632397

**Authors:** Marius Pachitariu, Lin Zhong, Alexa Gracias, Amanda Minisi, Crystall Lopez, Carsen Stringer

## Abstract

Artificial neural networks learn faster if they are initialized well. Good initializations can generate high-dimensional macroscopic dynamics with long timescales. It is not known if biological neural networks have similar properties. Here we show that the eigenvalue spectrum and dynamical properties of large-scale neural recordings in mice (two-photon and electrophysiology) are similar to those produced by linear dynamics governed by a random symmetric matrix that is critically normalized. An exception was hippocampal area CA1: population activity in this area resembled an efficient, uncorrelated neural code, which may be optimized for information storage capacity. Global emergent activity modes persisted in simulations with sparse, clustered or spatial connectivity. We hypothesize that the spontaneous neural activity reflects a critical initialization of whole-brain neural circuits that is optimized for learning time-dependent tasks.

## Introduction

The parameters of deep neural networks are typically initialized with independent random numbers chosen from standard distributions such as uniform or Gaussian [1]. Good initializations can substantially accelerate learning and lead to better final models [2, 3]. Commonly used initializations scale the amplitudes of the weight matrix by the inverse square root of the number of in-units, out-units or a combination of these [1, 4]. More complex initialization schemes are rarely tested (but see [5–7]), and the principles by which initialization helps are not well-studied. A better understanding of initialization may lead to better initialization schemes for modern, complex models such as transformers [8], state space models [9], and deep signal processing models [10].

Separately, in biological networks, we and others have found that intrinsically-generated neural activity contains macroscopic modes of coordination between neurons that extend across the entire mouse brain [11–14]. More activity variance was concentrated into the top dimensions of neural activity than would be expected for independently-firing neurons. At the same time, there was no low-dimensional cutoff of variance concentration, and the variance scaled as a power-law of the eigenmode number [11]. There are no mechanistic models yet that can explain this scaling of variance, but macroscopic variability in general has been hypothesized to arise from neural network dynamics operating in either a critical or chaotic regime [15–20]. We follow previous work to assume that the interactions in a high-dimensional neural network can be approximated by random matrices [21–25]. In other words, we assume that random matrices can summarize the combined effect of a large number of interactions in the network. We use basic properties of symmetric random matrices such as the semicircle law to explain the emergence of macroscopic patterns in neural networks [26].

Furthermore, we postulate a relation between parameter initialization and the intrinsic activity in biological networks, measured at rest or during spontaneous behaviors. Such spontaneous activity may reflect the initialization rule of the biological network, and may provide the scaffolding on top of which learning can occur. Supporting this hypothesis, we find that random matrix dynamics with stochastic inputs generate a variance spectrum that decays as a power-law, similar to the neural data. The exponent of this power-law was 0.7-0.85 across recordings, which is similar to the exponents generated by a symmetric random matrix (∼0.7) but different from those generated by non-symmetric random matrices ∼1.25. The theory predicts that systems with symmetric dynamics have no complex, rotational eigenmodes, and that the timescales of the eigenmodes correlates with their variances, two properties that also held in the neural data. These properties were independently validated in large-scale neural recordings from a single brain area (visual cortex / two-photon calcium imaging) as well as in recordings distributed over the entire brain (eight-probe Neuropixels / electrophysiology). In contrast, large-scale neural recordings in the hippocampus of passive animals had very different dynamics, mostly devoid of macroscopic structure.

## Results

### Random matrix dynamics generate macroscopic structure

Our initial modeling goal was to reproduce the power-law scaling of variance across modes of neural population activity [11]. We make the simplifying assumption that during spontaneous activity, the nonlinear network dynamics can be approximated as linear dynamics around a stationary point. In addition, we assume that each model unit (which could be a neuron or a group of neurons) generates independent stochastic variation, such as may be observed for example in the Poisson-like firing to external stimuli. When the stochastic inputs are Gaussian, this model describes a stochastic Ornstein-Uhlenbeck (O-U) process [19, 28, 29]. However, the results below do not require Gaussianity of the noise, only independence across units and across time. The interaction matrix *A* contains independent random, positive numbers uniformly distributed, representing the excitatory interactions between units (Figure 1a). To stabilize the dynamics, we subtract the mean of this matrix, which in the brain could be implemented with global inhibitory feedback (see Methods, Figure 1b). The results below still hold when the interactions are drawn from other distributions (see Figure S1).

**Figure 1:**
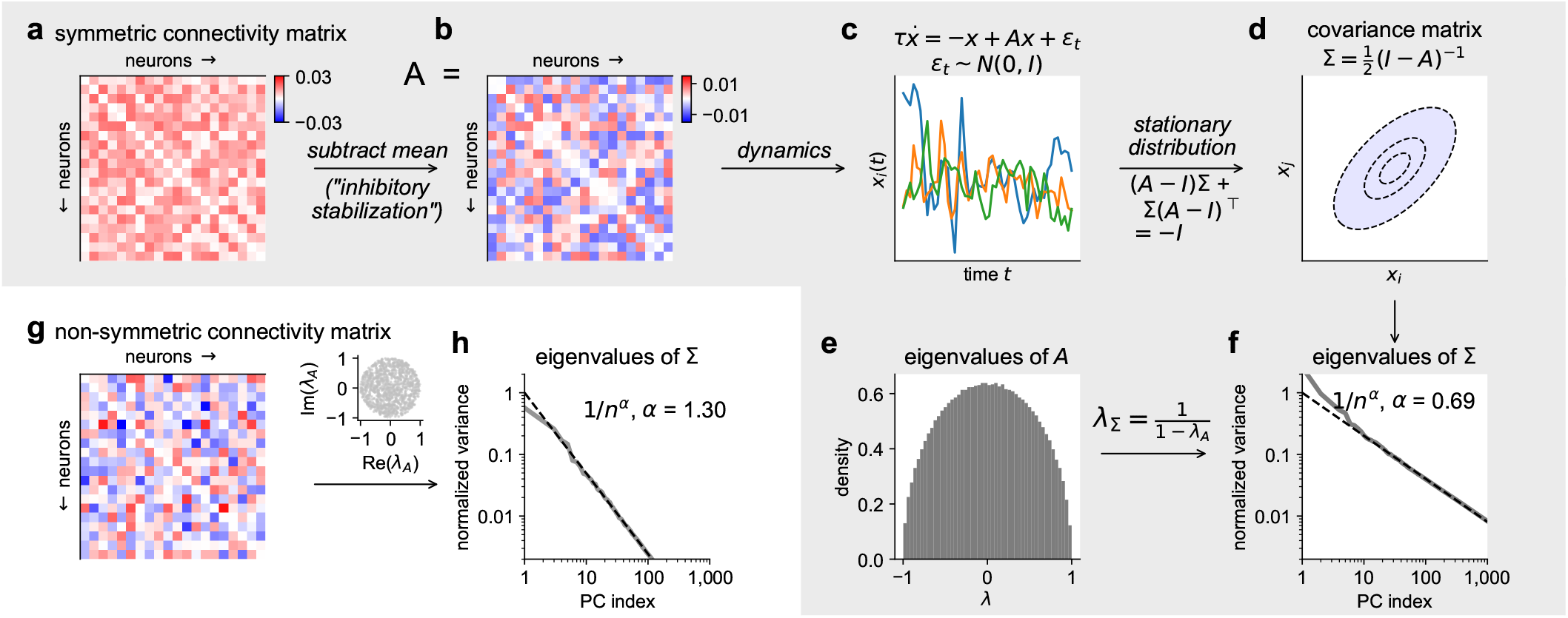
Dynamical system with random connectivity produces power-law covariance structure. **a**, Symmetric connectivity matrix *A*, with entries drawn from a uniform random distribution. **b**, Connectivity matrix from **a** with the mean of the matrix subtracted, representing global inhibition. **c**, Example neural activities from dynamical system with connectivity *A*. **d**, Covariance matrix Σ derived from the Lyapunov equation for the stationary distribution. **e**, Eigenvalues of random symmetric matrix *A* follow the semicircle law [26]. **f**, Sorted eigenvalues of covariance matrix Σ follow a power-law. **g**, Same as in **a**, but connectivity now is non-symmetric. Right inset: Eigenvalues of random asymmetric matrix *A* follow the circular law [27]. **h**, Eigenvalues of covariance matrix Σ follow a power-law with a faster decay than symmetric case.

When the interaction matrix *A* is symmetric, the covariance of the resulting multi-dimensional activity can be calculated directly from *A* via the Lyapunov equation [30], and its eigenvalues can be related to the eigenvalues of *A* (Figure 1c-d). We further scale *A* to have a spectral radius (i.e. largest eigenvalue) of 1 or close to 1 (see Methods). We call such matrices critically normalized. In this case, it can be shown both mathematically and numerically that the eigenvalues of the covariance decay as a power-law with exponent ∼ 2/3 (Figure 1e-f, Figure S2).

The case for non-symmetric interactions *A* proceeds similarly and results in a power-law of variances with approximate exponent 1.25 (Figure 1g-h, Figure S2). We could not obtain a direct mathematical estimate in this case and leave this as an open problem. When the interaction matrix is only partially symmetric, the eigenvalues decay as a power-law with intermediate exponent between 2/3 and 1.25 (Figure S2). Thus, critically-normalized matrices with varying degrees of symmetry can model the covariance spectrum of high-dimensional real-world datasets, which we have previously observed to follow power-law decays with varying exponents [34]. We focus here exclusively on the spontaneous activity observed in large-scale neural recordings from the mouse brain.

We note that some of our modeling assumptions are similar to [19] but our quantitative predictions (∼2/3 power-law for symmetric matrices) are very different from theirs (∼4/3 power-law). These are quite different predictions: in a 2/3 power-law for a population of 10,000 neurons, ~1,500 dimensions are required to account for 50% of the variance, while in a 4/3 power-law, 3 dimensions are sufficient.

The discrepancy is due to the theory in [19] being developed for very large binning windows, which may account for very slow timescales of neural activity but not the faster timescales that may be more relevant to neural computations. We found empirically that windows as large as 10 seconds are needed to reach a power-law of ∼ 4/3 in the simulations (Figure S3).

### Neural recordings match the structure of symmetric random matrix dynamics

We have previously estimated the power-law decay of spontaneous population activity in V1 and obtained exponents of 1-1.2 [11, 35], which would be in the middle of the range between symmetric and non-symmetric connectivity in the model. However, these estimates were upper bounds, because a test set was used to measure variances, and because the variances were estimated in either 1.2 or 0.3 second bins in transgenic mice expressing GCaMP6s, a sensor which attenuates higher frequencies in the neural activity. To obtain better estimates of the eigenvalue spectrum of the neural activity in small bins, we recorded at ∼22Hz from V1 in transgenic jGCaMP8s mice (6,433 - 10,595 ROIs), thus taking advantage of the much faster dynamics of jGCaMP8s (Figure 2a). In addition, we removed the train/test split from shared variance components analysis (SVCA), an eigenvalue estimation method (see Methods), and showed that this provides more accurate estimates in simulations (Figure S4). We also performed the eigenvalue estimation in brainwide neural recordings with 8 simultaneous Neuropixels probes (1,716 - 2,914 single-units, Figure 2b), which we also binned at 22Hz [11, 36]. Finally, we ran the same analyses on large-scale datasets from hippocampal area CA1 in GCaMP6f transgenic mice recorded at 22Hz (3,981 - 8,519 ROIs, Figure 2c). In all cases, mice were head-fixed in complete darkness, and performed spontaneous behaviors, and for all analyses based on two-photon (2p) calcium imaging, we used spike-deconvolved data (see Methods).

**Figure 2:**
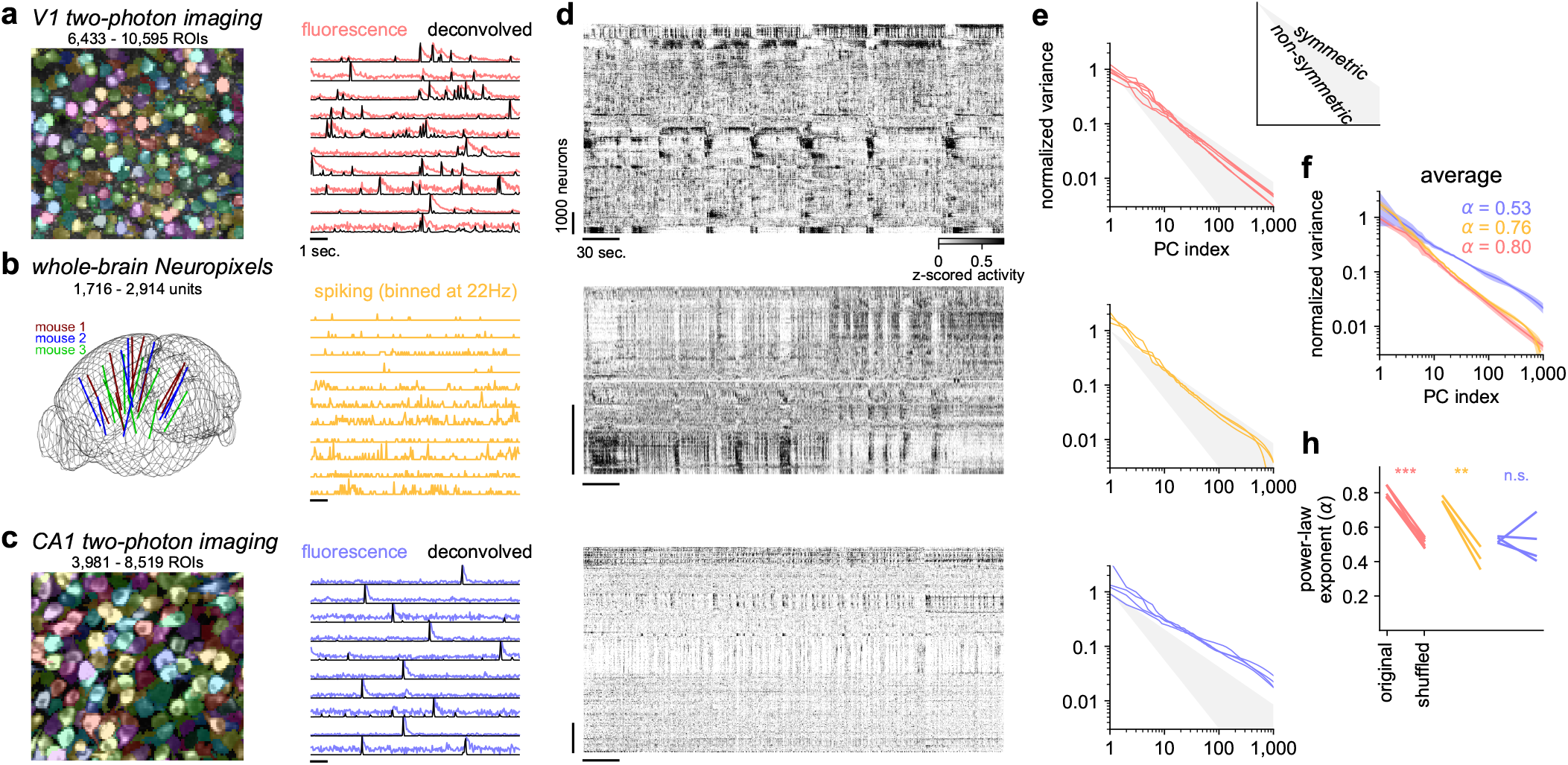
Power-law correlation structure in neural recordings. **a**, Two-photon calcium imaging in mouse V1, at 22Hz. Left: maximum projection image overlaid with units found. Right: Fluorescence traces of example neurons with deconvolved traces overlaid. **b**, Whole-brain Neuropixels recordings [31, 32], from [11]. Left: Locations of electrodes in Allen CCF reference [33]. Right: Firing from example neurons, binned at 22Hz. **c**, Same as **a**, from hippocampal CA1, at 22Hz. **d**, Rastermap [34] of all neurons from each recording displayed in **a**-**c. e**, Eigenvalue spectrum from each dataset. Shaded area bounded by power-law exponents from simulations with symmetric and asymmetric matrices. **f**, Average eigenvalue spectra across datasets. **h**, Power-law exponents of the decay of the eigenvalue spectra, for original datasets versus the neurons shuffled in time.

We first visualized the population recordings using Rastermap, a method for reordering neurons in a raster plot so that neurons with similar activity patterns are placed next to each other (Figure 2d) [34]. Qualitatively, the cortical 2p recordings and the brainwide ephys recordings displayed macroscopic coordination in neural firing, which was mostly absent in the CA1 recordings. The variance spectra of this activity decayed with power-law exponents in the range of 0.7-0.85 for both the V1 2p recordings and the brainwide ephys recordings, close to the estimates expected from the stochastic dynamics of a random symmetric matrix (Figure 2e-h). These exponents were substantially reduced after shuffling in time the activity of individual neurons (Figure 2h). In contrast, the hippocampal CA1 population activity had a variance spectrum that decayed much slower, with exponents in the range of 0.4-0.5, and which were not substantially altered by single-neuron shuffling (Figure 2e-h). Thus, the CA1 spontaneous activity was consistent with a model of nearly independent neural firing, which could be obtained for example from random matrix dynamics with a non-critical, over-normalized interaction matrix.

The model also predicts that the variance of the principal components should covary with their intrinsic timescales (see Methods). To verify the prediction in the data, we computed the auto-correlograms (ACGs) of the principal components of each recording (Figure 3a). We took care to estimate the ACGs in a manner that takes the noise level of each component into account (see Methods). In addition, we smoothed the ephys dataset in time in order to find the eigenvectors, but reordered these based on their raw, non-smoothed variance. This step was necessary due to the relatively fewer neurons and much shorter recording times in the ephys dataset, both of which increase the estimation noise of the eigenvectors. Across all datasets, we found that principal components with more variance had slower temporal dynamics, as predicted by the theory (Figure 3b).

**Figure 3:**
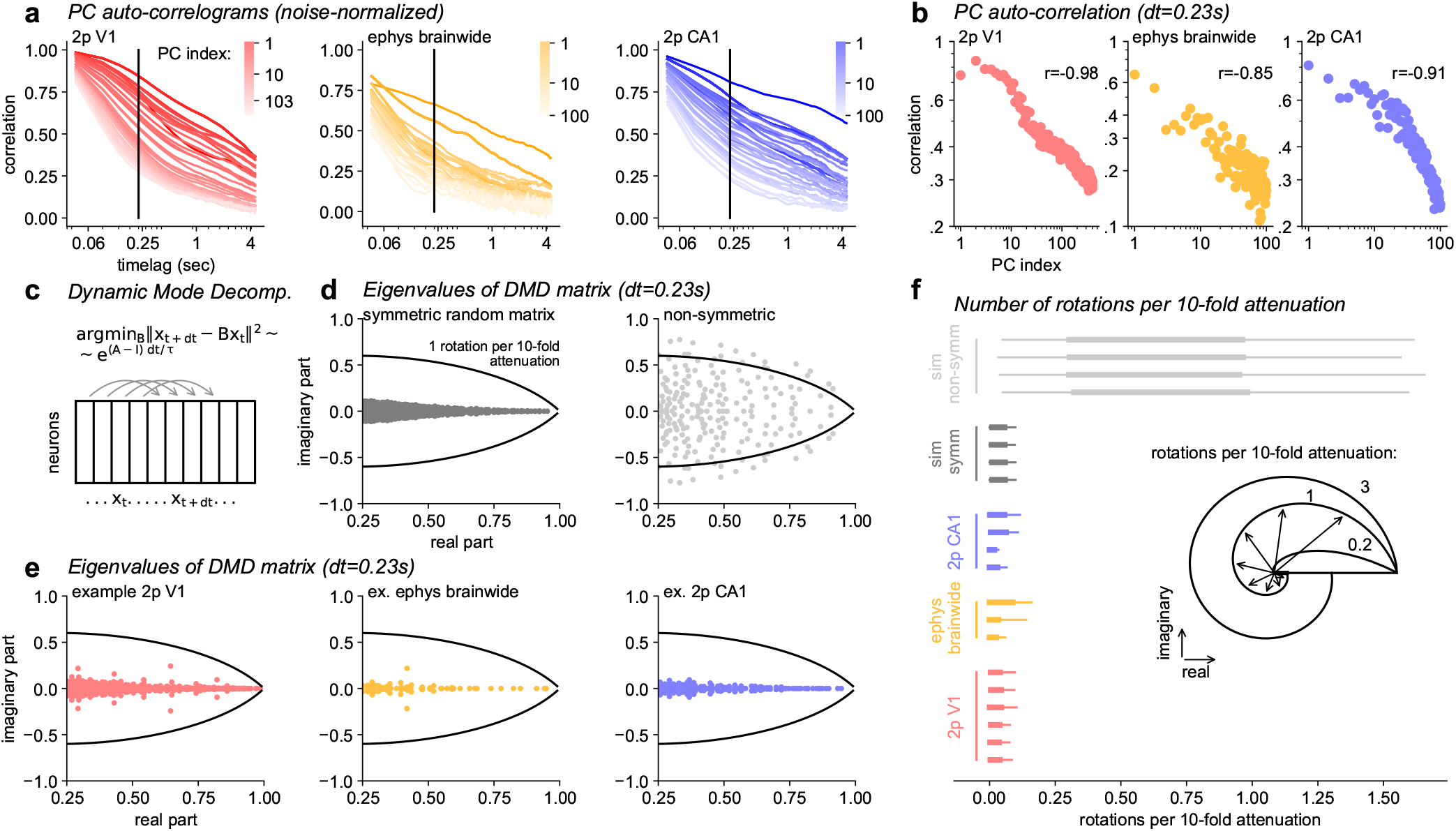
Dynamical properties of neural macroscopic dynamics. **a**, The estimated auto-correlograms of the neural PCs, averaged across all datasets of the same type. **b**, Average auto-correlation at a timelag of ∼0.23sec. **c**, Schematic of dynamic mode decomposition. **d**, Eigenvalues of the estimated DMD matrices at a timelag of ∼0.23sec for simulations with symmetric (left) and non-symmetric (right) random matrices. **e**, Same as **d** for example real datasets. **f**, Estimated number of rotations per 10-fold attenuation for the complex eigenvalues estimated from DMD, shown per dataset. Thick and thin lines indicate 25-75% and 5-95% ranges respectively.

Next, we investigated a key difference between symmetric and non-symmetric dynamics: the complexity of the eigenvalues of their dynamics. Symmetric matrices have real eigenvalues, while non-symmetric random matrices have eigenvalues distributed on a disk in the complex plane [27]. Symmetric, stable matrices produce relaxation dynamics, while non-symmetric random matrices produce substantial rotational dynamics. The Rastermap visualization gave us the first indication that the spontaneous neural activity does not contain rotational dynamics because there was no consistent drift or sequential activity across the neural subpopulations. To directly quantify this effect, we turned to time-lagged Dynamic Mode Decomposition (DMD) (Figure 3c), a popular method for identifying dynamical effects in sequential data [38, 39]. For linear dynamical systems, the eigenvalues of the DMD matrix are directly related to the eigenvalues of the linear dynamics matrix. In particular, the eigenvalues of the DMD matrices for symmetric random dynamics have nearly zero complex parts, unlike those from non-symmetric random dynamics (Figure 3d). Estimated from the data, the DMD matrices all had near-zero complex parts (Figure 3e). We quantified this with the number of full rotations per 10-fold attenuation of the magnitude of the complex eigenvector projection (Figure 3f). Only the simulated, non-symmetric dynamics reached levels of rotation that affect the dynamics significantly.

### Macroscopic dynamics persist under structured connectivity

So far, we have found the neural data to match the structure of dynamics with a critically-normalized, symmetric random matrix. This model however ignores many structural properties of real neuronal circuits. For example, connections between neurons are sparse, they depend on distance in tissue, and they also depend on cell types. In this section, we show that if at least a small fraction of the connections are global, then global activity modes still emerge, with the same power-law distribution of eigenvalues. We focus on symmetric matrices.

Sparse, symmetric random matrices also follow the Wigner semicircle law when the sparsity is not very high [40]. Thus, the dynamics of the resulting systems retain the same 2/3 power-law scaling of variance across activity modes. Empirically, we found that the connection probability needed to be at least 0.4% in a network of 10,000 units to leave the variance spectrum unchanged (Figure 4a). Similar properties apply to clustered and spatially-structured stochastic connectivity, as long as each connection is drawn from a mean-zero distribution (see Methods). In simulations, the variance spectrum remained unchanged as long as the global connection probability was at least 1% of the local probability (Figure 4b,c).

**Figure 4:**
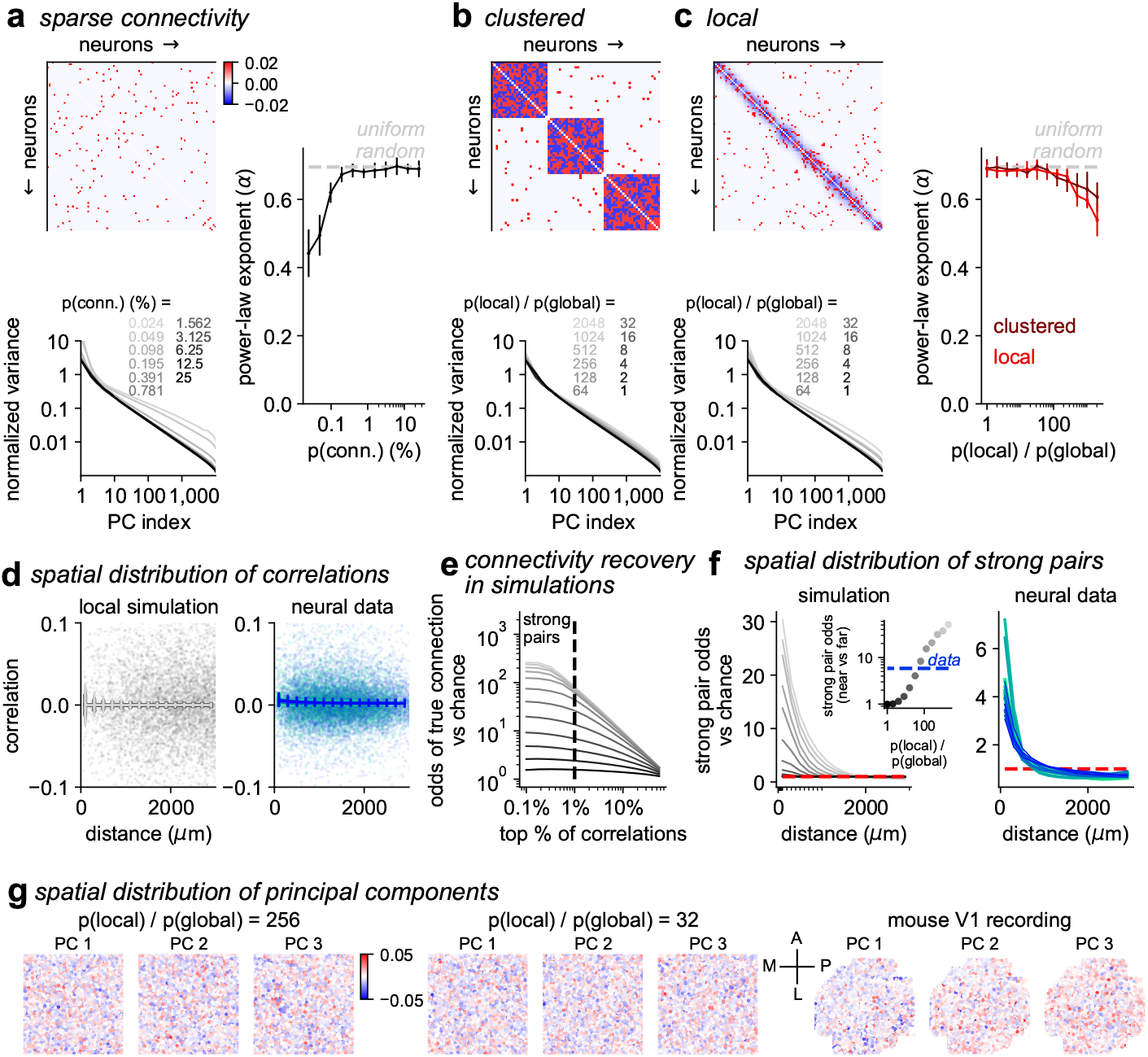
Persistence of global activity modes under biophysical connectivity patterns. **abc**, Top: Connectivity matrices with symmetric sparse / clustered / localized weights. Inhibition is set in proportion to the probability of connection for each weight. Bottom: variance decay of principal components for simulations with different levels of sparsity / clustering / localization. Right: fitted power-law exponents for these conditions (n=10 simulations per condition, error bars are standard deviation). Dashed line indicates the 0.69 value from Figure 1. **d**, Scatter plot of pairwise correlations vs distance in either: (left) a simulation with local connectivity, or (right) mesoscopic, single-unit recordings from [37]. Mean for each simulation/recording as a solid line, error bars are standard deviation across neuron pairs. **e**, Strong pairs (top 1% correlated pairs) are likely to be true connections across a wide range of sparse connectivity as in **a. f**, Dependence of strong pair odds with distance in either: (left) simulations with local connectivity or (right) recordings. Inset of left plot shows monotonic relation between the strong pair odds at near vs far distances (which can also be estimated from data) and the true ratio of local and global probability of connection. **g**, Top principal components are global in both recordings and simulations, even for very large bias towards local connectivity.

We investigated the spatial model further, since we could potentially match it to properties of spatially-sampled neural recordings from a previous study [35, 41]. The pairwise correlations across neurons in the simulation did not have significant distance-dependence, similar to the data (Figure 4d). Thus, average correlations cannot be used to infer the strength or length scale of spatial connectivity. We hypothesized that instead we could use the highest correlated pairs of neurons as a better match to connectivity. We define as “strong pairs” those pairs of units or neurons in the top 1% highest correlation. In simulations, the strong pairs had a much higher chance to be connected compared to chance (Figure 4e). The odds of a given pair being a strong pair varied strongly with distance, with a similar length scale as that used to simulate the data (Figure 4f). Further, the odds-ratio of strong pairs between near- and far-pairs was proportional to the ground truth probability ratio of local and global connections (Figure 4f). We observed a similar relation with distance of strong pair likelihoods in the data (Figure 4f, previously reported in [42]). Thus, strong pairs are enriched at short distances, and that may be a reflection of a higher connection probability. Finally, we note that in both data and in simulations with strong local connectivity, the top principal components are always global (Figure 4g).

## Discussion

The modeling results we presented are quite general. They hold with arbitrary distributions of the independent noise, with almost arbitrary distributions of the independent connectivity, and when various forms of structure are present in the connectivity matrix, such as low-rank or spatial structure. Across all these modeling choices, the substantial difference between symmetric and non-symmetric systems holds (2/3 vs ∼1.25 power-law exponent), which corresponds to a dramatic difference in effective dimensionality. The 0.75-0.8 power-law exponent in the neural recordings may therefore indicate that a high-dimensional code is preferable for neural computations, and that it is achieved through critically-normalized, symmetric interactions. Symmetric interactions are widespread in the brain: brain areas [43] and pairs of neurons [44] are often reciprocally connected. The critical normalization needed to generate these dynamics can be achieved in a self-tuned way as previously suggested in other contexts [45]. For example, an initially unstable system can be scaled down, by pruning or rescaling connections, until it is stable [46], through various mechanisms [47, 48].

Models with random connectivity and nonlinear dynamics have a long history in computational modeling [49–51]. These modeling efforts typically exploit the (near-) chaotic dynamics in such networks to perform nonlinear computations that require memory. Recent deep learning methods show that even models with linear dynamics can perform such tasks as long as they are deep [9, 52]. This led us to hypothesize that spontaneous activity in the mouse brain reflects the initialization of a brainwide neural network that may provide ideal conditions for computations that require dynamics and memory. There already exists evidence that this scaffold is used to represent motor states [11], and that laboratory tasks trigger a brainwide cascade of neural activity similar to the patterns observed in spontaneous activity [12, 53–55]. Perhaps all the learning that needs to happen in such tasks is on the feedforward connections from sensory inputs to the brainwide dynamical reservoir.

## Acknowledgments

This research was funded by the Howard Hughes Medical Institute at the Janelia Research Campus. We thank Gabriela Michel for GP5.17 animals, Caiying Guo for generating jGCaMP8s transgenic animals, Sara Barnes and other Vivarium staff for animal husbandry, Dan Flickinger for microscope maintenance, Jon Arnold, Alex Sohn, and Samuel Charles Jager for rig mechanical maintenance. We thank Juan Alvaro Gallego and Tosif Ahamed for discussions.

## Data availability

Data will be made available upon publication in a journal.

## Code availability

Code will be made available upon publication in a journal.

## Methods

All experimental procedures were conducted according to IACUC at HHMI Janelia. Data analysis and simulations were performed in python using pytorch and numpy, and figures were made using matplotlib and jupyter-notebooks [56–60].

### Data acquisition

#### Animals

All experimental procedures were conducted according to IACUC, ethics approval received from the IACUC board at HHMI Janelia Research Campus. We performed 6 recordings in cortex in 6 mice bred to express bred to express jGCaMP8s in excitatory neurons: TetO-jGCaMP8s x Camk2a-tTA mice (available as JAX 037717 and JAX 007004). We also performed 4 recordings in hippocampal CA1 in 4 mice bred to express GCaMP6f in excitatory neurons: Thy1-GCaMP6f GP5.17 mice (available as RRID:IMSR JAX:025393) [61]. These mice were male and female, and ranged from 2 to 12 months of age. Mice were housed in reverse light cycle, and were pair-housed with their siblings before and after surgery. Holding rooms are set to a temperature of 70°F +/- 2°F, and humidity of 50%rH +/- 20%.

#### Surgical procedures

Surgeries were performed in adult mice (P35–P125) following procedures outlined in [62] and [63]. In brief, mice were anesthetized with Isoflurane while a craniotomy was performed. Marcaine (no more than 8 mg/kg) was injected subcutaneously beneath the incision area, and warmed fluids + 5% dextrose and Buprenorphine 0.1 mg/kg (systemic analgesic) were administered subcutaneously along with Dexamethasone 7 mg/kg via intramuscular route. In the canula implants, the same total Dexamethasone dose was administered tapered over three days: 4mg/kg on the first day, 2mg/kg on second day and 1mg/kg on third day.

For the visual cortical windows, measurements were taken to determine bregma-lambda distance and location of a 4 mm circular window over V1 Cortex, as far lateral and caudal as possible without compromising the stability of the implant. A 4+5 mm double window was placed into the craniotomy so that the 4mm window replaced the previously removed bone piece and the 5mm window lay over the edge of the bone. For the hippocampal windows, the craniotomy was centered at 1.8 mm AP and 2.0 mm ML from bregma. Cortex was aspirated and a 3 mm glass coverslip attached to a stainless-steel was implanted over the dorsal CA1 region. CA1 surgeries were similar to [63].

After surgery, Ketoprofen 5mg/kg was administered subcutaneously and the animal allowed to recover on heat. The mice were monitored for pain or distress and Ketoprofen 5mg/kg was administered for 2 days following surgery.

#### Imaging acquisition

We used a custom-built 2-photon mesoscope [64] to record neural activity, and ScanImage [65] for data acquisition. We used a custom online Z-correction module (now in ScanImage), to correct for Z and XY drift online during the recording. As described in [62], for the visual area and hippocampal recordings, we used an upgrade of the mesoscope that allowed us to approximately double the number of recorded neurons using temporal multiplexing [66].

The mice were free to run on a styrofoam cylinder. Mice were acclimatized to running on the ball for several sessions before imaging, and one mouse was trained on a virtual reality task for two weeks prior to the recording. The field of view was selected such that large numbers of neurons could be observed, with clear calcium transients. Recordings were performed for 100-150 minutes at a rate of 22Hz. Recordings from [35, 41] were acquired at a rate of 3Hz.

#### Processing of calcium imaging data

Calcium imaging data was processed using the Suite2p toolbox (v0.9.3) [67], available at www.github.com/MouseLand/suite2p. Suite2p performs motion correction, ROI detection, neuropil correction, and spike deconvolution as described elsewhere [11]. We used a neuropil subtraction coefficient of 1.0. For the 22Hz recordings, we used all ROIs output by Suite2p above an SNR threshold of 0.3, which included dendritic processes, to increase the number of units recorded. The SNR for the activity trace *x* for each ROI was defined as

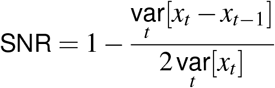

(similar to [68]). 61±16% (mean±s.d.) ROIs had an SNR greater than 0.3, resulting in a range of 3,981 - 10,595 ROIs across recordings.

We improved the spike deconvolution here by utilizing the latest version of Suite2p, which will be described in an upcoming manuscript. Our approach was similar to [68], where a neural network is used to predict the ground truth spikes from the noisy convolved traces. Unlike [68] we trained the model on a large number of simulations single-neuron spiking activity convolved with GCaMP-like dynamics and we used a Unet predictive model [69] with a style-vector to capture temporal context independently for each deconvolved trace [70]. We verified that the deconvolved traces better capture neural activity on real data with less noise by evaluating the responses to visual stimuli presented at known times.

#### Neuropixels recordings and processing

As described in [11], eight Neuropixels electrode arrays were used to record simultaneously from up to 3,000 neurons across the brain in three mice [31]. On the day of recording, mice were briefly anesthetized with isoflurane while eight small craniotomies were made with a dental drill. After several hours of recovery, mice were head-fixed in the IBL task setup: seated in a plastic tube with their forepaws on a wheel, surrounded by three computer screens in a light-isolated enclosure [53, 71]. The electrodes were advanced slowly (∼10 *µ*m/sec) to their final depth (4 or 5 mm deep), and allowed to settle for ∼15 minutes before recording. During the spontaneous part of the recording, the computer screens were black. Data were preprocessed by re-referencing to the common median across all channels [72].

These recordings were re-processed with Kilosort4 (v4.0.14), with default settings [73]. The Kilosort4 spike sorting found 1,756, 2,837 and 2,962 neurons defined as “good”, with refractory violation rate < 0.2 (default), from the 3 recordings. We excluded neurons with a firing rate of less than 0.01Hz during the spontaneous recording period, resulting in 1,716, 2,787 and 2,914 neurons in total in each recording. The spikes were binned at a rate of 22Hz, the same acquisition rate as the calcium imaging. The period of spontaneous activity in each recording was 22 to 42 minutes long (only this period was used for analyses).

### Data analysis

We normalized the neural activity in order to avoid fitting single neuron statistics. We z-scored the activity of each neuron so that the mean activity of each neuron is 0 and its standard deviation is 1. We ran Rastermap on the recordings with 100 clusters and 128 principal components, and visualized the sorted activity using 20 neurons per bin [34].

#### Eigenspectrum estimation

We estimated the eigenvalues using the covariance between two halves of neurons from the recording. This is to avoid contaminating the eigenspectrum with single neuron noise produced from the recording methods and from Poisson variability. We divided the recordings in half spatially, using a checkerboard of size 50*µ*m for the calcium imaging recordings, and using sections of 40*µ*m (8 contacts) on each Neuropixels probe in the electrophysiological recordings. We did not additionally split the data into training and testing timepoints, as we found this inflated the power-law exponent estimates (Figure S4) [11].

We estimated the power-law exponent of the eigenspectrum decay using a weighted linear regression in log space from rank 10 to 500, with weights as the inverse of the log of the rank [74]. The eigenvalue spectra are normalized by the value of the linear regression fit at rank 1.

#### Estimating PC timescales from data

To estimate the timescales of the principal components, we must take into account the noise. A naive estimation of the auto-correlograms would find that later PCs have smaller timelagged correlations, but that could simply be due to these PCs having lower SNR overall, and thus all their timelagged correlations would be lower. Instead, we take a similar approach to that from SVCA: we split the data into two random subsets of neurons using a checkerboard grid with size 50*µ*m, and we calculate the components with an SVD on the covariance between the two subsets of neurons. The resulting left and right singular vectors were used to project test data, and the correlograms were computed between the projections of one set of neurons and the other. The resulting estimates of the PC auto-correlograms were then normalized to 1 at a timelag 0.

#### Estimating rotational components from data

When the dynamics matrix *A* is not symmetric, its eigenvalues are complex, and the covariance of the multi-dimensional data is no longer directly related to these eigenvalues. Thus, to directly evaluate the complexity or rotational aspect of the dynamics, we cannot rely on the principal components alone. Instead, we fit linear predictive models that predict the neural population vector at time *t* + *dt* from the population vector at time *t*. Such models are typically referred to as dynamical mode decomposition (DMD), with the small modification that we use a timelag *dt* = 0.23sec instead of the more common *dt* = 1 time sample. The reason to use a timelag is to make estimation of the rotational modes more robust, and in particular to avoid the potential influence of short timescale artifacts arising from the deconvolution of the calcium imaging data.

To estimate DMD, we first reduced each dataset to 1,000 dimensions via PCA. We then used ridge regression with a penalty of 0.1 to predict *X*_*t*+*dt*_ from *X*_*t*_ in the reduced PCA space. Thus *X*_*t*+*dt*_ ≈*BX*_*t*_, with *B* a square matrix mapping PCs to PCs. As usual, the neural activity was z-scored on a per-neuron basis before applying PCA, but it was not re-normalized afterwards. Since *PCA* is an orthonormal projection, the eigenvalues of *B* are the same as would be expected in the full neuronal space, other than estimation errors. In the model, the matrix exponential describes the relation between *X*_*t*+*dt*_ and *X*_*t*_, regardless of whether *A* is symmetric or not:

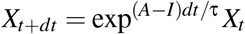

Thus, the DMD matrix *B* we obtained from data is an estimate of exp^(*A*−*I*)*dt*/τ^, at least in the simulations. Looking at the complexity of the eigenvalues of *B* can thus indicate whether the dynamics are rotational or not. Note that the eigenvalues λ′ of exp^(*A*−*I*)*dt*/τ^ are related to the eigenvalues λ of *A* by λ′ = exp^(λ−1)*dt*/τ^. Thus the higher the complex part of λ, the higher the complex part of λ′. We can also now see why taking a larger *dt* is beneficial: when *dt* is very small relative to the timescale of the dynamics, the eigenvalues λ′ approach 1, making it difficult to estimate their rotational component. The relation λ′ = exp^(λ−1)*dt*/τ^ could inverted to obtain estimates of λ. However, this multi-step process is likely to contain a lot of estimation error, and we preferred instead to directly compare the estimated distributions of λ′ from data to those from appropriately matched simulations. In particular, we compute the number *n*_*rot*10_ of rotations per 10-fold attenuation of the complex eigenvector λ′:

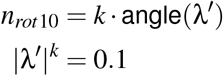

Thus:

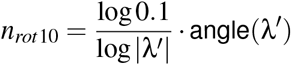

We used τ = 50ms in the simulations with symmetric matrices in order to approximately match the timescales of the data. Longer or shorter τ in the simulations would simply contract all estimated eigenvalues towards 0, but otherwise leave the number of rotations per 10-fold attenuation unchanged.

#### Local correlation structure

In Figure 4, we computed the correlations using the recording sampling rate (3Hz). For the simulations, we used the correlation matrix derived from the eigenvectors. For all the analyses we excluded neuron pairs within 20 *µ*m of each other. The pairwise correlations were binned in 200 *µ*m bins (Figure 4d).

We defined the top 1% of correlations per neuron as the “strong pairs” for each neuron. We then computed the probability distribution of the strong pairs across spatial bins of 200 *µ*m. This distribution was normalized by the distribution of all other correlations across bins, producing the strong pair odds versus chance shown in Figure 4f. The strong pair odds, near versus far, was the ratio of the first bin of this curve (within 200*µ*m) to the average of the last four bins of this curve (2200-3000*µ*m).

### Dynamical systems analysis

We assume that neural activity is governed by linear dynamics with independent stochastic inputs. In the case of normally-distributed inputs, this model becomes the familiar Ornstein-Uhlenbeck process, with connectivity *A* [28]:

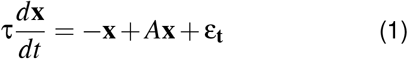

where the noise is a Wiener process.

The stationary distribution of the neural covariance matrix Σ is defined by the Lyapunov equation [30]:

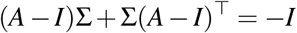

When *A* is symmetric, the solution is given by

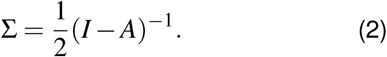

For the asymmetric case, the solution must be calculated numerically [75] (see also [76] for an example of solving for *A*).

For the symmetric case, we can derive the decay of the eigenvalue spectrum directly, under the assumption that *A* is a random symmetric matrix with an eigenspectrum distribution of a semicircle from -1 to 1. We assume here that the number of units is large enough to treat the eigenvalue distribution as an exact semicircle distribution and ignore finite size effects. The rank *n* of eigenvalue 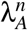 of *A* is defined by the integral of the semicircle distribution density 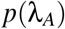 from 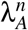 to 1. We scale *p*((λ_*A*_)) to a maximum of 1 for convenience of the calculations. If we define θ as the angle subtended by 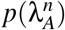 on the semicircle, we can use geometric arguments to show that:

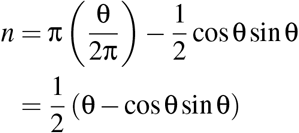

The eigenvalues of the covariance Σ, denoted by λ, are related to the eigenvalues λ^*A*^ of the connectivity matrix *A*:

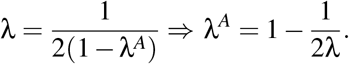

Thus, we have

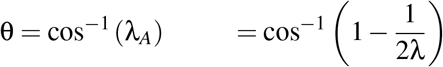

Plugging this into the equation for rank *n* and using 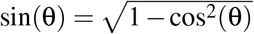 we have:

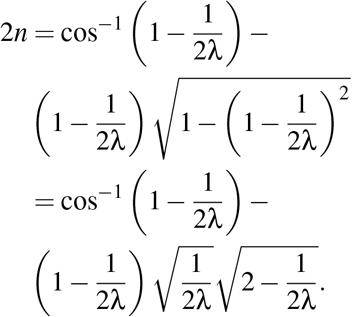

We next used Wolfram Alpha [77] to obtain the Taylor expansion for 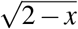 and Puiseux expansion for cos^−1^(1− *x*) where *x* is small. Keeping only terms in which *x* is raised to a power less than 2, gives

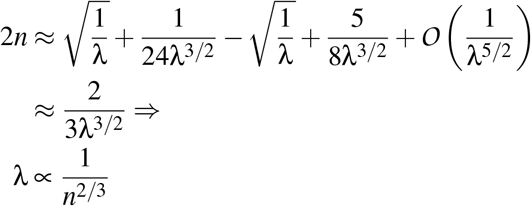

Thus, the eigenvalues decay approximately as a power-law with exponent 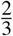, which is very close to the value we found in simulations.

As mentioned in the text, this is different from the value of 4/3 found in [19]. The discrepancy is due to [19] estimating the “long time window covariance”, which assumes that the data is binned in infinitely-long windows. The authors of [19] argue that the formula converges for short windows (>50 ms), which appears to use the single-neuron timescales as a reference. However, when the dynamical systems are close to critical, the emergent timescales are much longer, and thus a much longer window is needed to reach the stable state. Thus, the derivation of [19], while interesting, can only capture the infraslow timescales of neural activity and predicts a 4/3 power-law on a rank plot in this case.

We note that the 4/3 exponent, while not explicitly calculated there, can be easily derived from equation 16 in [19]:

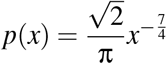

which integrated gives:

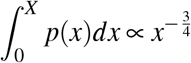

which results in a power-law exponent of 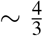 in a rank plot [19].

#### Relating timescales to eigenvalues

In addition to estimating the decay of variances of the principal components, we also want to evaluate dynamical, temporal properties of the data and relate them to the model. In the model, the timescales of the system are related to the eigenvalues of *A* and therefore to the eigenvalues of Σ, following from the matrix exponential solution to the Lyapunov equation:

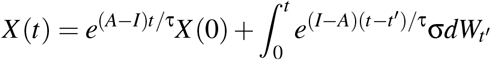

The second term on the right is a noise term that is independent of *X*(0). Using the SVD decomposition of *A* = *U* Λ*U*^*T*^, with Λ having the eigenvalues 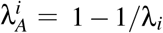 on the diagonal, the multidimensional system can be divided into independent scalar equations:

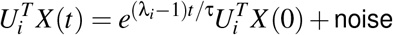

Thus, the PC component projections 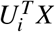 have an autocorrelation that decays like

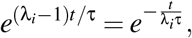

and thus the timescales of the PCs are monotonic with the amplitude of the eigenvalue λ_*i*_. For the purposes of the analysis in Figure 3, we need only observe the cross-correlation at timelag *t*, though its exact decay with λ_*i*_ cannot be predicted owing to the unknown single unit timescale τ.

### Simulations of dynamical systems

We simulated 10,000 neurons governed by the dynamics in Equation 1. For the dynamics simulations, we performed integration using the Euler-Maruyama method and used a step size of 2 ms and a timescale τ for each neuron of 50 ms using pytorch [57, 78]. The random noise was drawn from a Gaussian with a mean of zero and standard deviation of one for each neuron at each time step. We ran 200 simulations on a GPU in parallel, with random initial conditions drawn from a Gaussian with mean zero and standard deviation one, each consisting of 60,000 time steps, and discarded the first 4,000 time steps. To replicate the sampling rate in the data, we binned every 23 timepoints (∼22 Hz). We also z-scored the unit activities, as in the data. For testing the eigenspectrum estimation methods (Figure S4), we added Gaussian random noise (mean zero, standard deviation one) to each timepoint of the binned and z-scored simulated traces. We then smoothed the traces in time with a Gaussian of standard deviation of 1 timepoint (at 22Hz) to approximate the smoothing in the calcium imaging. We used only 10 out of the 200 randomly initialized simulations per random connectivity matrix to replicate the limited time in the neural recordings.

#### Dense connectivity

We drew the excitatory connections between neurons from a uniform random distribution from zero to two. We subtracted off the mean connectivity (one). We set the diagonal values of the matrix to zero. When this matrix is symmetric, its eigenspectrum distribution follows the semicircle law; for the non-symmetric case, it follows the circular law [79]. We divided the matrix by a scalar so that the largest real value of the eigenvalues of the matrix was 0.999, setting *A* so that it is critically normalized.

#### Sparse / varied connectivity

We created the sparse symmetric random matrices with random zero or one connections drawn from a Bernoulli with a mean varying from 2.4e-4 to 0.25 (Figure 4a). The mean of the Bernoulli was subtracted globally from the entire matrix, resulting in a matrix with zero mean connectivity. The diagonal of the matrix was set to zero. Random sparse symmetric matrices and graphs also follow the semicircle law, when the sparsity is not too high [40, 80–82].

We created the clustered symmetric random matrices by setting the Bernoulli mean to 0.5 for within-cluster (local) connectivity, and the mean out-of-cluster (global) connectivity in a range from 2.4e-4 to 0.5 (Figure 4b). This results in a ratio of the probability of local connection to the probability of global connection (p(local) / p(global)) ranging from 1 to 2048. Each cluster consisted of 500 neurons. The mean of the Bernoulli distribution for each entry was subtracted (0.5 within-cluster, smaller outside), resulting in mean zero connectivity across the matrix. The diagonal of the matrix was set to zero.

We created the locally-connected symmetric random matrices using as the Bernoulli mean an exponential decay function of Euclidean distances *d*_*ij*_ in *µ*m between neurons in the simulation (Figure 4c). The exponential function was defined as 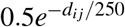. The neurons were placed randomly on a torus of size 8000 *µ*m by 8000 *µ*m. The minimum value of the mean of the Bernoulli was set to a value from 2.4e-4 to 0.5, resulting in a range of p(local) / p(global) from 1 to 2048. Again, the mean of the Bernoulli was subtracted from each entry of the matrix, and the diagonal of the matrix was set to zero. To quantify the recovery of true connectivity (Figure 4e), we compute the fraction of true positive connections with *p* % of top pairwise correlations per neuron, and then normalize by the average probability of positive connection in the simulation (chance).

As in the dense case, we divided each matrix by a scalar so that the largest eigenvalue of the matrix was 0.999. Because these connectivity matrices were symmetric, we directly computed the covariance matrix and the eigenspectrum from *A* using Equation 2.

---

**S1:**
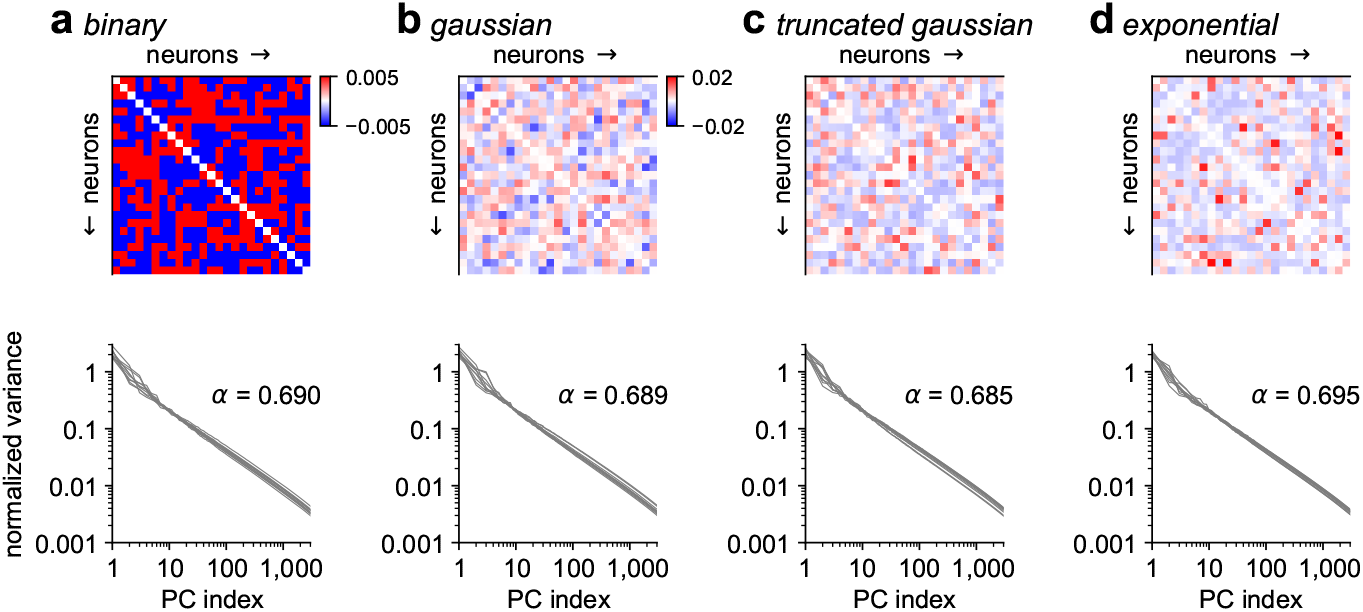
Simulations of dynamics from connectivity matrices with different probability distributions. **a-d** Top: connectivity matrices, in which independent connections are drawn from various distribution: Bernoulli, Gaussian, truncated Gaussian and exponential. In all cases, the mean of the distribution is subtracted off, and values are scaled to a spectral radius of 0.999. Bottom: Eigenspectra of covariance matrices from 10 simulations with random instantiations of the connectivity matrix, average α reported across the simulations.

**S2:**
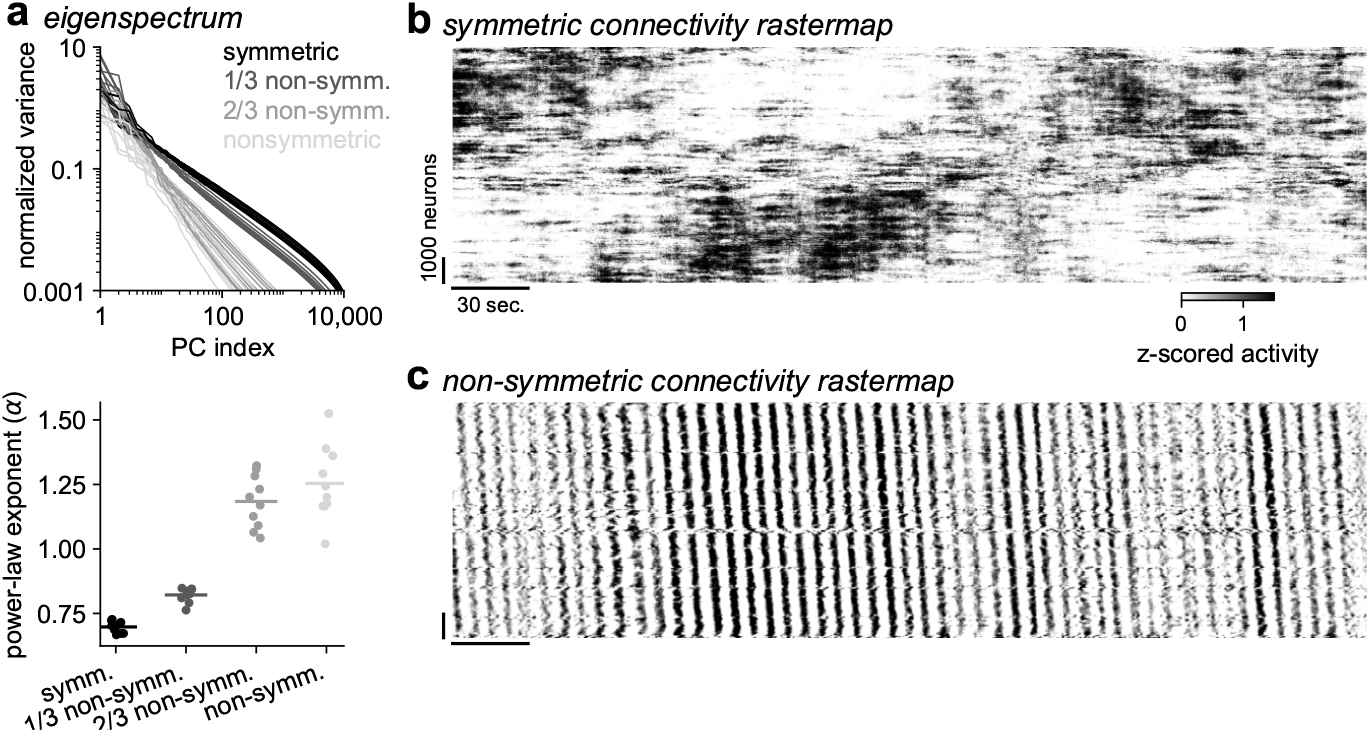
Simulations of dynamics from connectivity matrices with varying dynamics. **a**, Eigenspectra of covariance matrices from 10 simulations with connectivity defined by random symmetric matrices, random nonsymmetric matrices, and matrices with partial symmetry (1/3, 2/3). **b**, Rastermap of activity from an example simulation from a random symmetric connectivity matrix. **c**, Same as **b** for a random non-symmetric matrix.

**S3:**
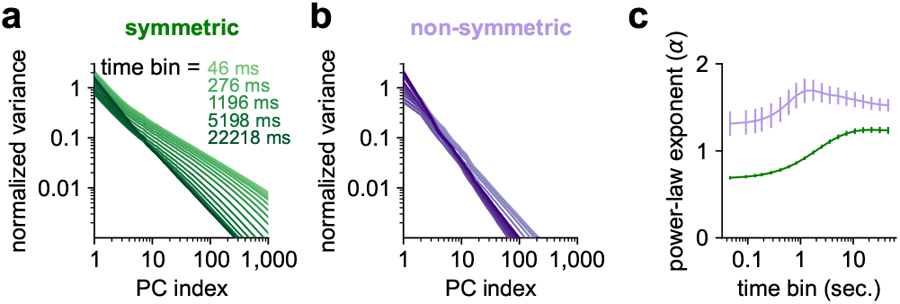
Effect of bin size on power-law exponent in simulations. **a**, Distribution of variance across principal components for simulations binned at different bin sizes, using a random symmetric, critically-normalized connection matrix. **b**, Same as **a** for non-symmetric connectivity matrices. **c**, Summary of fitted power-law exponents as a function of bin size for the simulations in **a** and **b**.

**S4:**
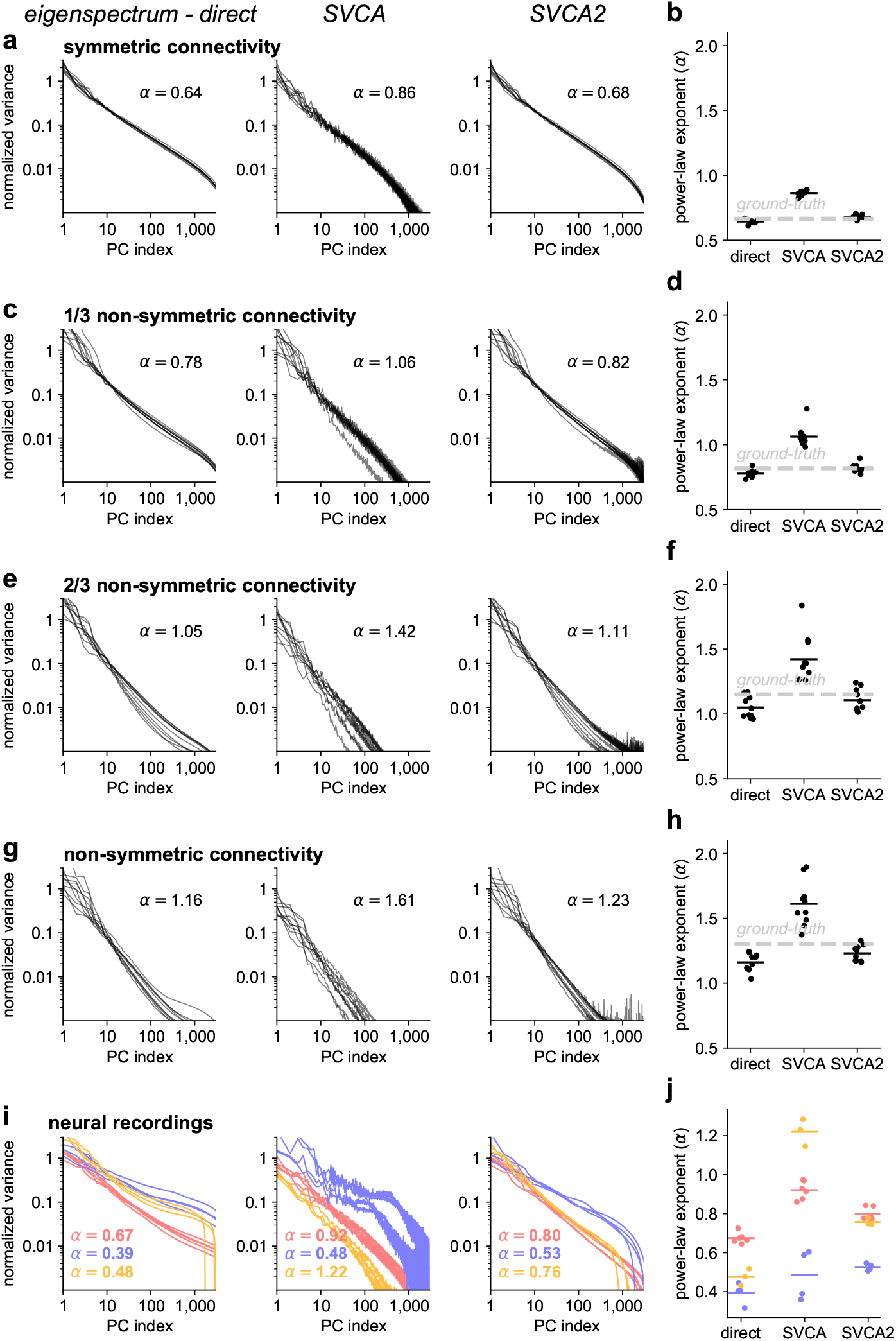
Eigenvalue estimation on simulations with added noise and smoothing, and on recordings. **aceg**, We added noise and smoothing to the unit activities of 10 simulations each for 4 conditions with various degrees of symmetry in the connectivity matrix (Figure S2). Left: Eigenspectrum of covariance of noisy and smoothed activity traces, estimated directly. Middle: Estimation of eigenspectrum using SVCA [11]. Right: Estimation of eigenspectrum using a modified version of SVCA without a train/test split in time (called “SVCA2”). **bdfh**, Power-law exponents for each estimation technique, with the mean power-law exponent from the ground-truth (no noise or smoothing) as a dashed line. **ij**, Similar to **ab** for real recordings, where the ground truth is not known.

